# Postmitotic nuclear pore assembly proceeds by radial dilation of small ER membrane openings

**DOI:** 10.1101/141150

**Authors:** Shotaro Otsuka, Anna M. Steyer, Martin Schorb, Jean-Karim Hériché, M. Julius Hossain, Suruchi Sethi, Moritz Kueblbeck, Yannick Schwab, Martin Beck, Jan Ellenberg

## Abstract

The nuclear envelope has to be reformed after mitosis to create viable daughter cells with closed nuclei. How membrane sealing of DNA and assembly of nuclear pore complexes (NPCs) are achieved and coordinated is poorly understood. Here, we reconstructed nuclear membrane topology and structure of assembling NPCs in a correlative three dimensional electron microscopy time-course of dividing human cells. Our quantitative ultrastructural analysis shows that nuclear membranes form from highly fenestrated ER sheets, whose shrinking holes are stabilized and then dilated into NPCs during inner ring complex assembly, forming thousands of transport channels within minutes. This mechanism is fundamentally different from interphase NPC assembly and explains how mitotic cells can rapidly establish a closed nuclear compartment while making it transport-competent at the same time.

## Introduction

The nuclear pore complex (NPC) is the largest non-polymeric protein complex in eukaryotic cells and composed of multiple copies of around 30 different proteins termed nucleoporins (Nups) ^1^. NPCs are the sole gates of macromolecular transport across the double membrane of the nuclear envelope (NE). In higher eukaryotes, NPCs and the NE disassemble at the beginning of mitosis and their rapid reformation during mitotic exit is essential for establishing a functional nucleus in the daughter cell ^2-4^.

The process of postmitotic assembly of the NPC and the nuclear membranes from mitotic ER has been studied *in vitro* using nuclei assembled in *Xenopus* egg extract and by live cell imaging using fluorescence microscopy. Several molecular players regulating the process have been identified, including inner nuclear membrane proteins, ER shaping proteins such as reticulons, nuclear pore components ELYS and Nup107−160 complex, nuclear transport receptors and Ran ^2-4^. In addition, kinetic observations of the bulk NPC formation across the NE has shown that postmitotic assembly proceeds by sequential addition of Nups in a clear temporal progression, that is almost identical between rodent and human cells ^5,6^.

Despite these important insights, the mechanism of NPC assembly after mitosis has remained unclear and is highly debated ^7-10^. Whether postmitotic NPC assembly is initiated in an already sealed NE and the NPC is inserted into this double membrane by a *de novo* fusion event^11,12^ similar to NPC assembly during interphase^13^, or if it starts already on the naked DNA and the membrane only later engulfs assembling NPC from the side ^14,15^, has remained unanswered. How thousands of NPCs can assemble within a few minutes without interfering with the rapid sealing of NEs during mitotic exit thus has remained mysterious. A major reason for this gap in our knowledge was that individual NPCs and ER topology are below the resolution of live cell fluorescence microscopy that is needed to capture the dynamic process of postmitotic nuclear assembly, precluding reliable and quantitative observation of NPC assembly and the sealing of NE membranes. Here, we combine live cell imaging with high resolution 3D electron microscopy to ultrastructurally reconstruct the dynamic process of postmitotic NPC and NE assembly.

## Results

### Nuclear membranes form from highly fenestrated ER sheets

To measure how nuclear membrane sealing around DNA relates to NPC formation in space and time, we reconstructed whole dividing human cells with a time resolution of approximately one minute after the beginning of mitotic chromosome segregation by correlating single cell live imaging with focused ion beam scanning electron microscopy (FIB-SEM) (Fig. 1a and **Supplementary** Fig. 1). Segmentation of membranes in close proximity to chromosomes showed that the layer of mitotic ER that contacts chromosomes exhibits a high degree of fenestrations (Fig. 1b,c and **Supplementary Movie 1**) as reported previously ^16^. At early times only about 10% of the chromosome surface was associated with ER, but starting at about 5 min after anaphase onset (AO), the surface of ER-chromosome contacts increased rapidly covering the chromosomes with newly formed NE within 2 min (Fig. 1b,d and **Supplementary Table 1**). Fine 3D segmentation of ER/NE membranes (**Supplementary Movie 2**) in the large volume EM data showed that at early times (up to 3.9 min) variably sized holes made up 43% of the surface of the ER sheets contacting chromosomes (Fig. 1c,e) and that 59% of these discontinuities displayed a diameter below 100 nm (Fig. 1f), i.e. on the order of NPCs. The degree of fenestration and hole size in the ER sheets contacting chromosomes decreased rapidly (Fig. 1c), with holes making up only 16% of the surface two minutes later and now 75% of them having a diameter below 100 nm (**Fig. e,f**; 6.3 min). This data demonstrates that the NE forms from highly fenestrated ER sheets that contain a very large number of discontinuities whose diameter shrinks as the ER-derived NE covers the chromosomes (Fig. 1d).

**Figure 1.**
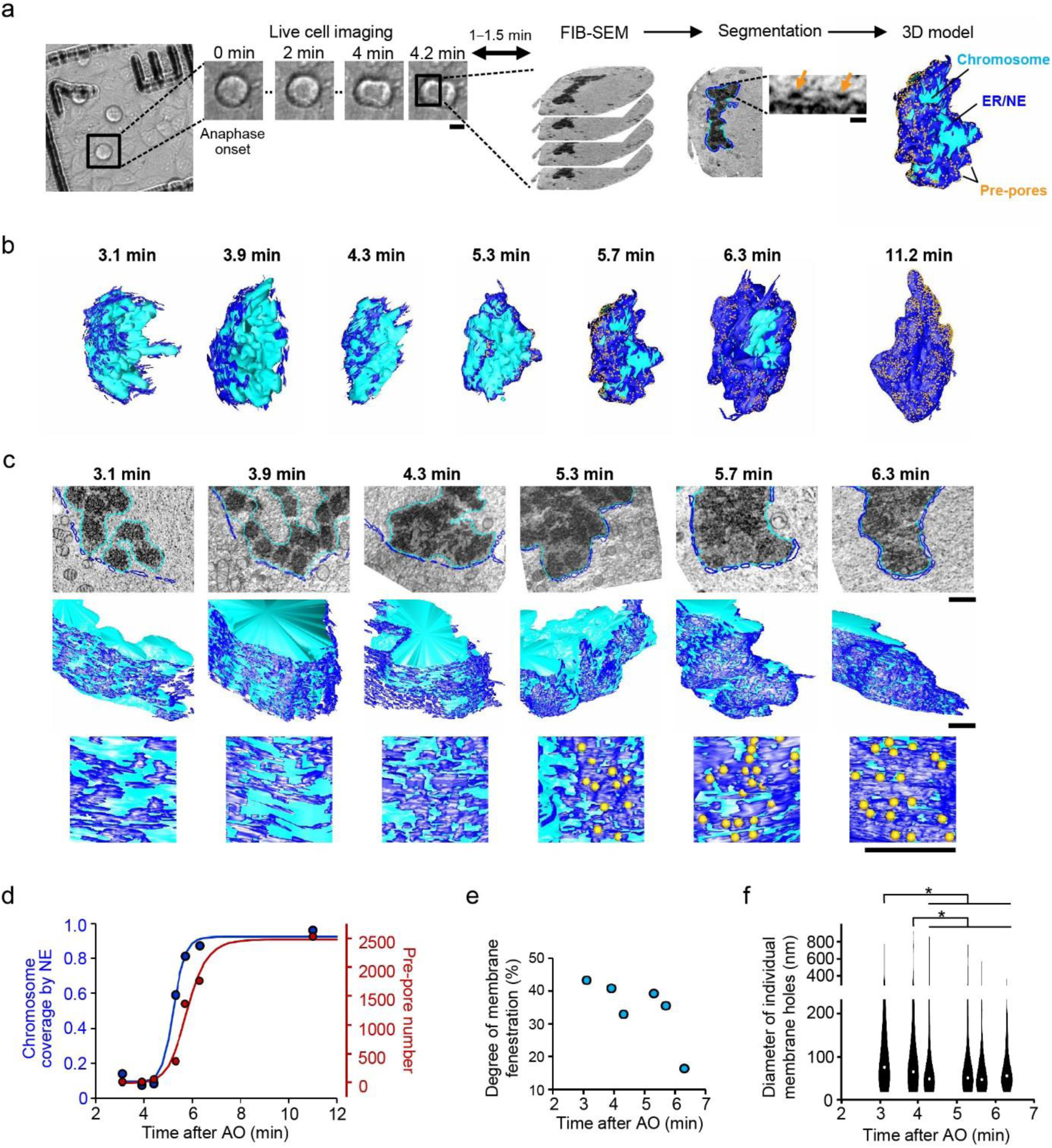
The nuclear envelope (NE) forms from highly fenestrated ER sheets. (**a**) Correlative live imaging with FIB-SEM. Cell division of HeLa cells was monitored every 0.2 min by light microscopy, and the time from anaphase onset (AO) to the time when the cells were frozen was measured for each cell. The data of 5.7 min cell is shown as an example. Dark red arrows indicate the pre-pores. The chromosome, the ER membranes in proximity to the chromosome, and pre-pores were segmented and shown as a model. Scale bar for LM, 10 μm; for EM 100 nm. (**b**) Models of entire nuclei at different times after AO. (**c**) Fine models for the selected parts of the nuclei. Examples for the segmentation, the model overview, and the enlarged view with pre-pores are shown in the upper, middle and lower panels, respectively. Scale bars, 1 μm. (**d**) Chromosome surface coverage by the NE and the pre-pore number in cells at different postmitotic times. t1/2 of the sigmoidal fit for NE sealing and pre-pore appearance were 5.2 and 5.8 min, respectively. (**e**) The ratio of free space to the membrane on the ER/NE sheet at indicated times. (**f**) The diameter of membrane holes. The violin plot is from 216, 188, 371, 289, 245, and 202 holes for 3.1, 3.9, 4.3, 5.3, 5.7, and 6.3 min, respectively. The median is depicted as a dot. *p<0.02; one-sided Mann-Whitney-Wilcoxon U test with Holm-Bonferroni correction for multiple testing.

### Coverage of chromosomes by nuclear membranes is closely linked to pre-pore formation

As ER fenestrae started to shrink significantly as early as 4.3 min after AO (Fig. 1f), many of the pore sized discontinuities started to contain electron dense material (Fig. 1a-c and **Supplementary Table 1**) and could therefore be classified as pre-pores. From their first appearance, the number of such pre-pores increased rapidly to 2000 with the local density of 15 pores/μm^2^ within only 3 min (Fig. 1d). Kinetic analysis of chromosome coverage by newly forming NEs and the appearance of pre-pores showed that both processes display sigmoidal kinetics and are closely linked in time with pre-pore appearance reaching its half-maximum within less than one minute after chromosome coverage (Fig. 1d).

### NPC assembly proceeds by dilation of small membrane holes

Knowing when exactly pre-pores start to form during NE formation, enabled us to examine the architecture of assembling NPCs at an even higher resolution. We performed correlative live imaging with electron tomography, in cells captured every 1−2 min after AO (Fig. 2a and **Supplementary Movie 3**), starting at 4.8 min when pre-pores first appear (Fig. 1d) until 15 min when NE formation is completed (Fig. 1b,d). Since NE sealing is delayed in the so called ecore-regions due to clearance of dense spindle microtubules ^4^ (Fig. 1b), we focused our analysis on the non-core regions of the NE (Fig. 2a). In a total of 27.8 μm^2^ reconstructed NE surface area, we identified 360 particles consistent with pre-pores (i.e. displaying a NE discontinuity containing regular electron dense material) captured at different times of postmitotic assembly (Fig. 2b, **Supplementary Fig. 2a** and **Supplementary Table 2**). At early times we also found 50 small NE discontinuities which were very similar to holes present in the ER not yet in touch with the chromosome surface (**Supplementary Fig. 3**). We classified these as small holes in NE or ER whose associated density is too low to be detectable as a distinct regular structure, although they might contain smaller amounts of proteins.

**Figure 2.**
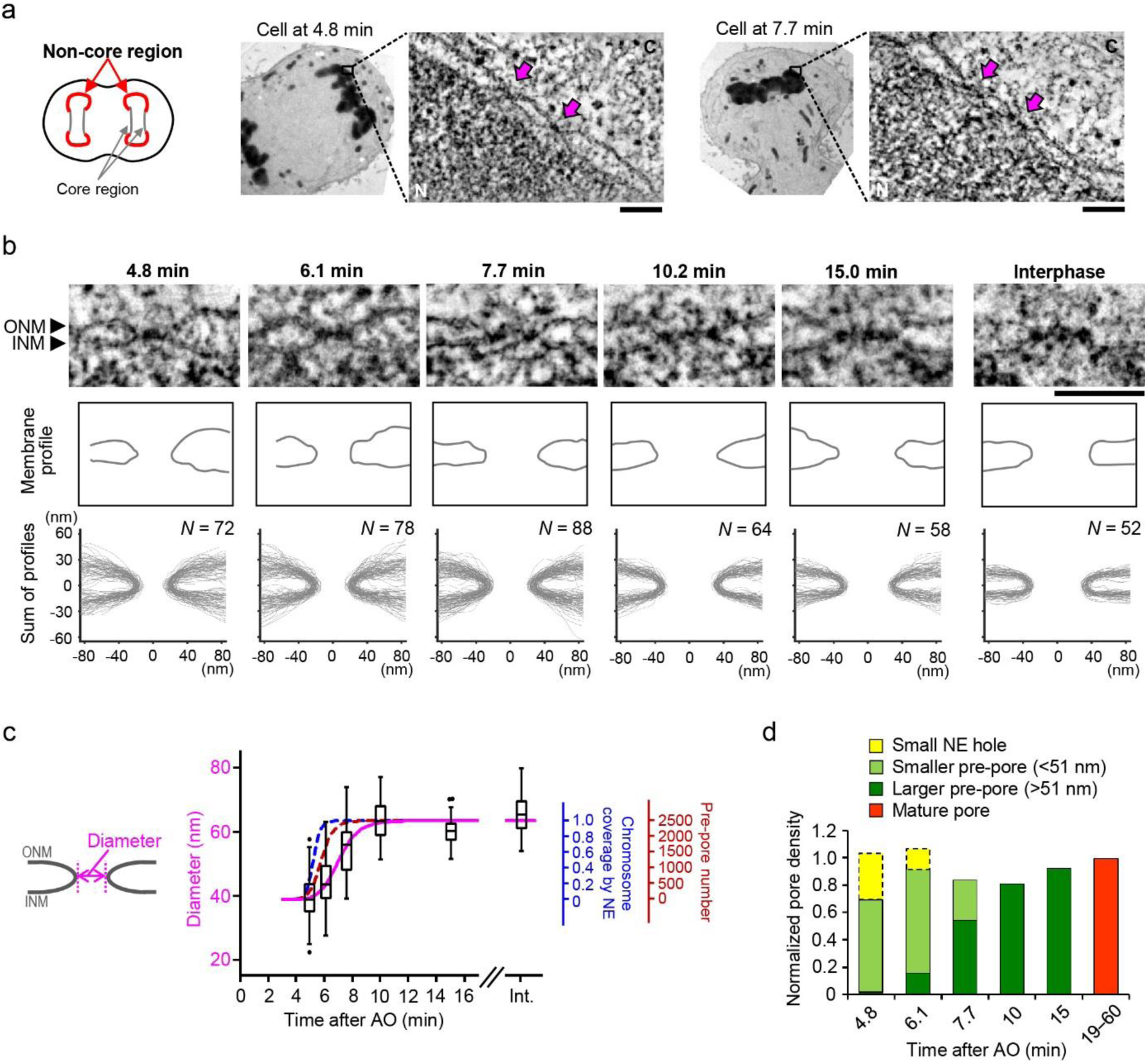
NPC assembly proceeds by dilation of small membrane holes. (**a**) Electron tomography of non-core regions of the NE (indicated in red in the left panel) in cells captured at different times after AO. Two examples are shown in the right panels. Enlarged images are tomographic slices of the non-core regions of cells at 4.8 and 7.7 min. Pre-pores are indicated by magenta arrows. C, cytoplasm; N, nucleoplasm. Scale bars, 100 nm. (**b**) Electron tomographic slices of pre - and mature pores at different times after AO. Scale bar, 100 nm. Membrane profiles of the individual pores (middle panels) and all the pores (bottom panels) at each time point are shown. ONM, outer nuclear membrane; INM, inner nuclear membrane. (**c**) Quantification of the pore diameter as indicated by a bidirectional arrow in the left panel. Center line, median; box limits, upper and lower quartiles; whiskers, 1.5x interquartile range; points, outliers. A magenta line shows a fitted sigmoid curve. Sigmoid curves of chromosome coverage by NE and pre-pore number in **Fig. 1d** are shown by dashed lines for comparison. (**d**) Normalized density of small NE holes, pre-pores smaller and larger than 51 nm, and mature pores at different times (see **Supplementary Fig. 2−4** for details).

We first focused our analysis on changes in NE topology. Tracing of the pre-pore membrane profiles in the 3D tomograms and their quantitative analysis revealed that pre-pore diameter increased rapidly from 39 nm (4.8 min after AO) to 63 nm (10.2 min) at which size they stabilized (Fig. 2b,c). The profile analysis also revealed other interesting NE topology changes (**Supplementary Fig. 2b,c**). The pre-pore dilation showed sigmoidal kinetics, reaching its half-maximum within 1.2 min after pre-pore appearance (Figs. 1d and 2c), predicting that it represents a maturation step into fully assembled NPCs. Detailed analysis of the distribution of pre-pore diameters at different postmitotic times allowed classification into two groups, smaller and larger pre-pores with a mean diameter of 42 and 62 nm, respectively (**Supplementary Fig. 4**). As predicted, the combined abundance of smaller and larger pre-pores matched the number of mature NPCs found after completion of nuclear reformation (Fig. 2d, **Supplementary Fig. 5** and **Supplementary Table 2**). Interestingly, at the beginning of pre-pore appearance (4.8 min), the slightly lower than expected density of smaller pre-pores was made up by the presence of similarly sized small NE holes, which disappeared at later times (Fig. 2d). Overall, this data indicates that pre-pores mature by membrane hole dilation into fully-assembled NPCs and that small NE holes are likely to be precursors of pre-pores that have not yet accumulated a significant amount of dense material.

### NPC assembly proceeds by centrifugal formation of a membrane associated ring

We next analyzed how the distribution of dense material inside pre-pores changes during their maturation. We first radially averaged all density inside the membrane hole in the NE plane of pre-pores (Fig. 3a). The change in mean radial intensity profiles over time showed that initially (4.8 min), pre-pores contained material in the center of the membrane hole; From 4.8 to 10 min, material progressively accumulated next to the membrane, resulting in a growing intensity peak that was close to the expanding wall of the membrane channel. After this peripheral accumulation of material, the center of the channel accumulated additional density from 10 min to interphase, leading to a second central peak in the radial profiles (Fig. 3a and **Supplementary Fig. 6**).

**Figure 3.**
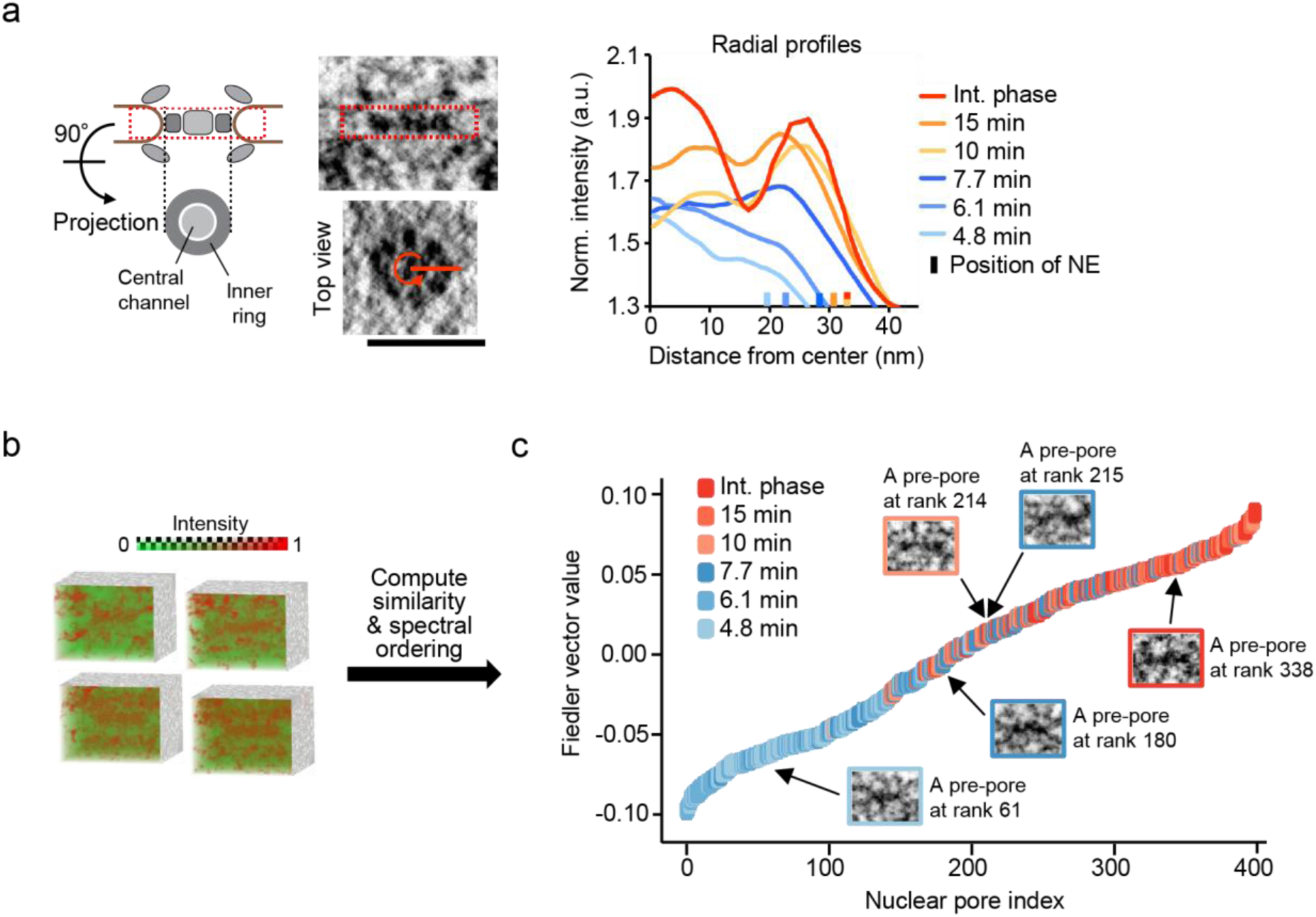
Intensity distribution inside the pre-pore membranes and unbiased classification of pre-pores from spectral ordering. (**a**) Radial profile analysis of pre-pores. The region indicated by red dashed box in the left panel was Z-projected and the radial intensity from the center of the pore was measured. Intensity was normalized to that of NE lumen. The mean radial intensities for each time point are shown (see also **Supplementary Fig. 6**). The positions of the membrane wall are indicated by short lines on the X-axis. Scale bar, 100 nm. (**b**) 3D intensity distribution of nuclear pores in the electron subtomograms. Four examples are shown. (**c**) Seriation of nuclear pores. Individual pre - and mature pores were ranked by the Fiedler vector value derived from the similarity of their 3D intensity distributions (see Methods for details). Individual nuclear pores are color-coded according to their time after AO. Five pores at indicated ranks of are shown as examples.

### Subtomogram averaging revealed a clear structural maturation of pre-pores

To obtain better insight into the structural changes during NPC assembly, subtomogram averaging of single pre-pores in the same state of assembly is necessary. Since the distributions of membrane hole diameters and profiles of dense material indicated that individual pre-pores sampled at the same time-point can vary in structure (**Supplementary Figs. 4** and **6**), we ordered them independent of time based on structural similarity using spectral seriation (Fig. 3b,c). Such spectral ordering of pre - and mature pores overall recapitulated their temporal sampling during anaphase, with pores at early (4.8 and 6.1 min), middle (7.7 min), and late (10, 15 min and interphase) time points ranked together (Fig. 3c and **Supplementary Fig. 7**), showing that postmitotic NPC assembly is indeed a progressive process. Based on their structural similarity, we partitioned pores into five assembly states (**Supplementary Fig. 7**) and performed subtomogram averaging. The averages revealed a striking progression of structural changes during postmitotic NPC assembly (Fig. 4a). Early and smaller pre-pores (cluster 1), exhibited dense material in the center of a narrow membrane gap, which subsequently shifted centrifugally towards the membrane (cluster 2) and then dilated into a clear peripheral ring with a first sign of the 8-fold rotational symmetry of the NPC inner ring complex (cluster 3). Inner ring complex formation was then completed with clear 8-fold symmetry (cluster 4), which was followed by maturation of the central channel density (cluster 5, Fig. 4a). Below the double nuclear membranes, density consistent with the nuclear ring was present from the beginning (cluster 1), whereas cytoplasmic ring-like density above the NE appeared only later (cluster 3, Fig. 4a). The same order of inner ring formation and dilation on top of an early assembled nucleoplasmic ring, followed by cytoplasmic ring assembly and central channel maturation, was also observed if pre-pores were clustered only according to experimental time (**Supplementary Fig. 8a,b**), showing that the pore maturation process is largely synchronous. Analysis of the increase in inner ring complex intensity over time in time-clustered averages furthermore showed that its sigmoidal rise coincides with the process of membrane dilation (**Supplementary Fig. 8c**), suggesting that inner ring complex self-assembly could drive pore dilation.

**Figure 4.**
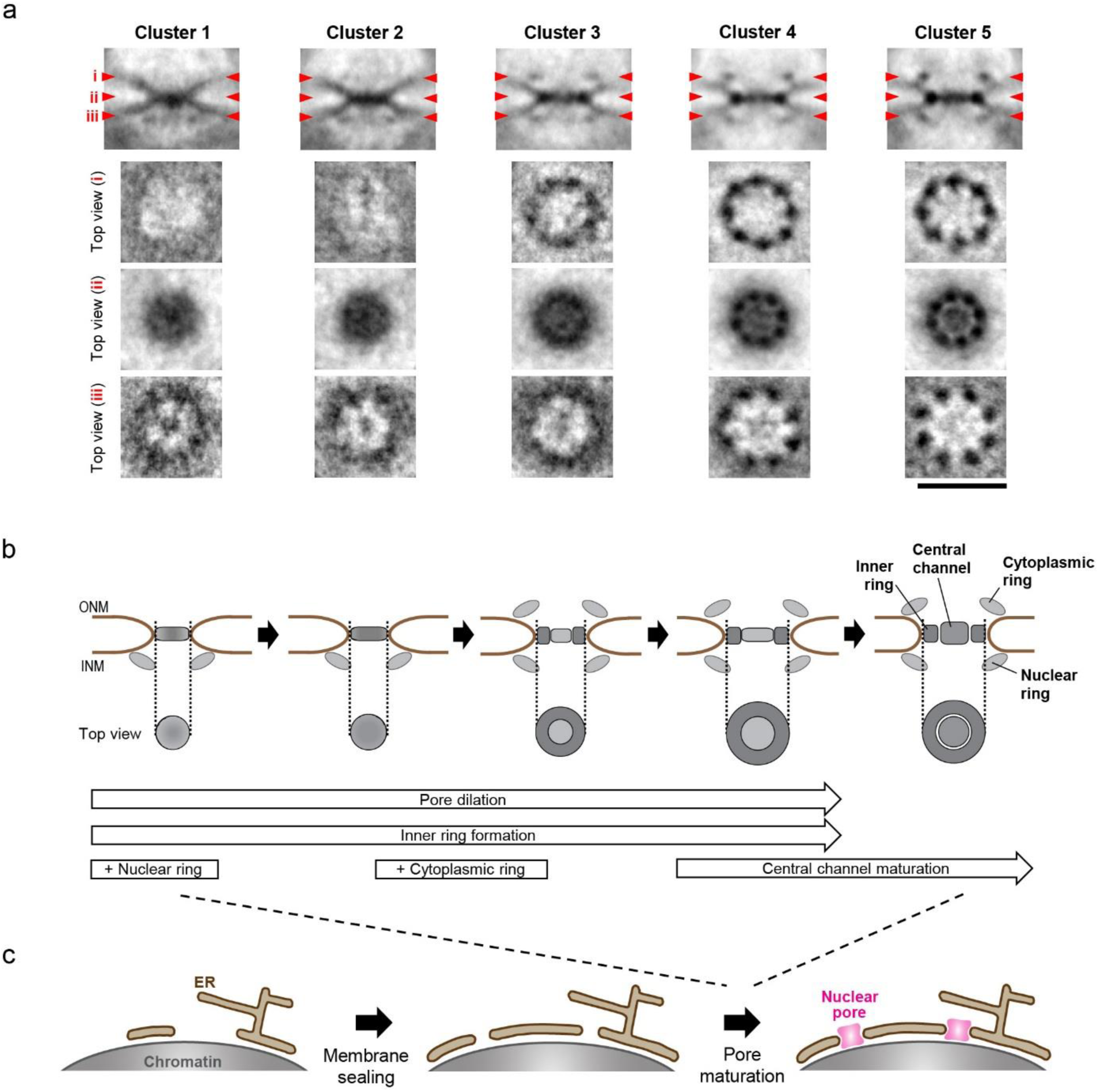
Structural maturation of pre-pores revealed by subtomogram averaging and postmitotic NE/NPC assembly model. (**a**) Electron tomographic slices of averaged nuclear pores in each cluster. No symmetry was imposed in averaging. The averaged images are from 61, 85, 69, 108, and 76 pores in cluster 1−5. Red arrowheads i, ii, and iii on side-view images indicate the locations of the planes which are inclined at 90° in top views i, ii, and iii. Scale bar, 100 nm. (**b**) A schematic model of postmitotic nuclear pore assembly. The initial pre-pore consists of some dense material in the center of the pore as well as the nuclear ring. During pore dilation, the inner ring and the cytoplasmic ring form. The pore dilates even further, and then the central channel is filled with additional materials. (**c**) A schematic model of NE assembly. The NE forms form a highly-fenestrated ER sheet that contains sufficient discontinuities for NPC assembly. The holes on the NE shrink during the membrane sealing, and the NPC assembly is initiated on the pre-existing holes on the NE.

The structural progression we observed is consistent with previous observations in live cells that nuclear/cytoplasmic ring components (Nup107 and Nup133) start to accumulate in the NE early, followed by the inner ring component Nup93 ^5,6^. We revisited these observations, by creating genome-edited cells in which nucleoporins are endogenously tagged with GFP to avoid effects of non-physiological expression levels ^13^. High time resolution live cell imaging and kinetic analysis of postmitotic NE accumulation showed that Nup205, a component of the inner ring complex, is incorporated after Nup107, a component of the nuclear and cytoplasmic ring complexes (**Supplementary Figs. 9** and **10**), consistent with our previous observations ^5,6^. Interestingly, the concentration of Nup107 on the NE had reached its half-maximum already at 6 min after AO when pre-pores contain only the nuclear ring, and increased further until 8 min when the cytoplasmic ring appears (**Supplementary Figs. 8b** and **10b**). The live cell kinetics support the notion that the observed nuclear ring in early pre-pores contains Nup107-160 subcomplex members, and the long accumulation of Nup107 is furthermore overall consistent with the EM finding that the nuclear ring assembles first followed by the later appearance of the cytoplasmic ring. Finally, the live cell accumulation of Nup205 shows similar kinetics to the increase in inner ring complex density over time observed by EM (**Supplementary Fig. 10c**), consistent with the idea that Nup205 contributes to inner ring assembly observed in EM. It should be noted that postmitotic NPC assembly is not perfectly synchronous at the single pore level as seen by the structural variability of pre-pores at each time point (**Supplementary Figs. 4** and **6**), and therefore the live bulk measurement of NE protein accumulation cannot provide the precise molecular assembly choreography of single pores.

## Discussion

How thousands of NPCs assemble into the reforming NE during mitosis exit has remained unclear due to the resolution limitation of conventional microscopy typically used to observe this dynamic process. Pioneering studies that used *in vitro* assembled nuclei with *Xenopus* egg extract and isolated nuclei from *Drosophila* embryos could unfortunately not establish the physiological nature of assembling NEs and NPCs, because the native membrane structure was disrupted during the egg extract preparation and nuclear isolation, and the observation was done on the outer surface of nuclei as they used scanning electron microscopy ^17,18^. Our temporally-staged ultrastructural analysis for the first time resolved postmitotic NE and NPC assembly in situ in intact human cells at nanometer resolution, enabling us to formulate a data-driven model of its mechanism (Fig. 4b,c). The fact that ER sheets that form the NE contain a sufficient number of small discontinuities for NPC assembly already at early time points (3.1, 3.9, and 4.3 min; Fig. 1f), and that we did not observe holes smaller than 20 nm in the NE although our resolution is sufficient to resolve them (Fig. 2 and **Supplementary Fig. 4**), strongly suggests that postmitotic NPC assembly starts in pre-existing small NE openings rather than by *de novo* fusion into already sealed double membranes. While our EM data cannot exclude that individual nucleoporins, that are not yet assembled into higher order structures, are already present in these small holes, they are morphologically clearly different from pre-pores and we cannot detect reproducibly positioned or ring-like densities in them (Fig. 2 and **Supplementary Fig. 3**). Since the first pre-pores containing such densities have a similar small size of around 40 nm diameter (Fig. 2b,c), it is likely that the ER/NE hole shrinkage is stalled and stabilized by protein accumulation in the center of the membrane hole. At later time points, pre-pores then dilate the membrane hole to normal NPC size of about 60 nm during inner ring complex formation, the cytoplasmic ring assembles and the central channel matures (Fig. 4b,c). The fact that an inner ring component Nup205 starts to be incorporated at 6 min after AO in live cells (**Supplementary Fig. 10**), argues that the density in the center of pre-pores at 4.8 min may not be explained by Nup205 and according to our earlier work also not much Nup93 ^5^. The most likely candidate protein to explain this early density would therefore by exclusion be Nup155, which forms the innermost layer of the inner ring complex ^19^ and has indeed been shown to be required for recruiting other inner ring components Nups205, 188 and 93 ^20,21^.

We have previously shown that *de novo* assembly of NPCs into intact nuclei during interphase proceeds via inside-out extrusion of the INM and fusion with the ONM ^13^, which is fundamentally distinct from the dilation mechanism of pre-existing membrane holes during postmitotic assembly reported here. While interphase NPC assembly takes about 45 min and is sporadic and rare ^13^, the rapid radial dilation of small membrane holes concomitant with inner ring complex formation supports the assembly of ~2000 postmitotic NPCs in only 3 min during NE sealing after mitosis. One of the reasons for this very high efficiency could be that assembly into the holes of highly fenestrated mitotic ER sheets does not require a new membrane fusion at the assembly site that is needed for interphase NPC assembly ^13^ and could represent a rate-limiting step. In addition, the mitotic cell contains a high concentration of assembly ready NPC subcomplexes, that become permissive for assembly synchronously by the reduction in mitotic kinase activity ^22^, while an interphase cell has to synthesize nucleoporins for assembly. Combined, this could explain the high efficiency of postmitotic NPC assembly that is essential for the rapid establishment of functional nuclei to exit mitosis. Our finding that the NPC assembles via a fundamentally different mechanism in mitosis than in interphase provides the basis to dissect the key structural and molecular transitions and regulatory steps in the future.

## Methods

### Cell culture

Wildtype HeLa kyoto cells (RRID: CVCL_1922; kind gift from Prof. Narumiya in Kyoto University) were grown in Dulbecco’s Modified Eagle’s Medium (DMEM) containing 4.5 g/l D-glucose (Sigma Aldrich, St. Louis, MO) supplemented with 10% fetal calf serum (FCS), 2 mM l-glutamine, 1 mM sodium pyruvate, and 100 μg/ml penicillin and streptomycin. The mycoplasma contamination was examined by PCR every 2 or 3 months and was always negative. For correlative light–electron microscopy, cells were cultured on sapphire disks (3 mm diameter; Wohlwend GmbH, Sennwald, Switzerland), and for three-dimensional (3D) fluorescence time-lapse imaging, cells were grown on 2-well Lab-Tek Chambered Coverglass (Thermo Fisher Scientific, Waltham, MA).

### Sample preparation for electron microscopy

Cells on pattern-branded sapphire disks by carbon-coating were incubated in imaging medium (IM; CO2-independent medium without phenol red (Invitrogen), containing 20% FCS, 2 mM lglutamine, and 100 μg/ml penicillin and streptomycin), and the cell division was monitored every 12 sec by widefield microscopy (Axio Observer Z1; Carl Zeiss, Oberkochen, Germany) using 10 × 0.25 NA A-Plan or 20 × 0.4 NA Plan-Neofluar objective (Carl Zeiss) at 37°C in a microscope-body-enclosing incubator. When cells entered the cell-cycle stages of interest, they were immersed in IM containing 20% Ficoll (PM400; Sigma Aldrich) for protecting cells from freezing damage, and then instantly frozen using a high-pressure freezing machine (HPM 010; ABRA Fluid AG, Widnau, Switzerland). It took 1.0−1.5 min from the last time-lapse imaging until the high-pressure freezing, and the time lag was always recorded to precisely determine the duration after anaphase onset. For focused ion beam scanning electron microscopy (FIBSEM), cells were freeze-substituted into Durcupan resin (Sigma Aldrich) as follows: Frozen cells were incubated with 1.0% osmium tetroxide (OsO4), 0.1% UA, and 5% water in acetone at −90°C for 20–24 h. The temperature was raised to −30°C (5°C/hour), kept at −30°C for 3 h, and raised to 0°C (5°C/hour). Samples were then washed with acetone, infiltrated with increasing concentrations of Durcupan in acetone (25, 50 and 75%), embedded in 100% Durcupan and polymerized at 60°C for 4 days. For electron tomography, cells were freeze-substituted into Lowicryl HM20 resin (Polysciences Inc., Warrington, PA) as described previously ^13^, and sections of 300 nm thickness were cut with an ultramicrotome (Ultracut UCT; Leica, Wetzlar, Germany) and collected on copper–palladium slot grids (Science Services, München, Germany) coated with 1% Formvar (Plano GmbH, Wetzlar, Germany). The Durcupan resin is very hard and good for precise FIB-milling. Freeze substitution into Lowicryl resin provides finer structure information than the one into Durcupan resin with OsO4. Therefore we used Durcupan and Lowicryl resin for FIB-SEM and electron tomography, respectively.

### Preparation for FIB-SEM acquisition

After high-pressure freezing and freeze-substitution, the sapphire disk was removed with the cells remaining at the surface of the resin block. The block containing the cells was first coarsely trimmed down with a handsaw and then finely with razor blades to make the monolayer cells as parallel as possible to the block-face. To be able to target the cells of interest in the SEM, marks were branded on the surface of the resin disk using a pulsed 20 × 355-nm laser (DPSL-355/14, Rapp Optoelectronic, Hamburg, Germany) coupled in an Olympus CellR widefield microscope using a 20 × 0.7 NA UPlanApo (Olympus) objective with the program called Xcellence cell^frap. The branded pattern left by the carbon coating is visible at the LM. After laser branding of the same landmarks, it is visible on the surface of the sample by the SEM, which can then be used to find back the region of interest (ROI) for image acquisition in the FIB-SEM.

### FIB-SEM

The resin disk containing the cells was mounted on a conductive carbon sticker (12 mm, Plano GmbH) that was placed on SEM stubs (6 mm length, Agar Scientific, Stansted, UK). If necessary, the parallel alignment between the disk and the SEM stub was done using small wedges of polymerized Durcupan. To limit charging by the electron beam during SEM imaging, the samples were surrounded by silver paint and coated by gold sputtering for 180 sec at 30 mA in a sputter coater (Q150R S; Quorum Technologies, Laughton, UK). The samples were then introduced into the Auriga 60/Crossbeam 540 (Carl Zeiss) and positioned so that the sample was facing the SEM at an angle of 36° and the FIB at an angle of 54°. ATLAS 3D, part of Atlas5 software (Fibics, Ottawa, Canada), was used to prepare the sample for “Slice & View”. As a first step a protective platinum coat of about 1 μm was deposited using the FIB at 1 µA/700 pA current (for Auriga 60/Crossbeam 540) to protect the surface of the sample and ensure even milling ^23^. In some datasets tracking lines were milled into the platinum coat with 50 pA FIB current and filled with SiO2 using 1 nA/700 pA current. An additional layer of platinum was added to protect the marks. The ion beam was used to dig a trench in front of the ROI with 16 nA/15 nA to reach a depth of 30 μm. After these surface modifications, the image surface was polished at 2 or 4 nA/3 nA. For imaging, the FIB was set to 1 or 2 nA/1.5 nA current, and SEM imaging and FIB milling were performed simultaneously ^23^. The images were acquired at 1.5 kV with the energy selective backscattered (EsB) detector with a grid voltage of 1.1 kV and the analytical mode at 700 pA current (specific for Crossbeam 540). The pixel size in xy was set to be 5 nm and the FIB-milling was done every 8 nm. The dwell time and line average was set for the frame-time to be 1.5 min.

### Segmentation of chromosome, ER, nuclear envelope and nuclear pore

The acquired raw FIB-SEM data were aligned using TrackEM2 ^24^ in ImageJ (http://rsbweb.nih.gov/ij/). After alignment, the stack was cropped, gray levels were inverted and the images were smoothened with a mean filter (kernel size: 15 × 15 nm). For some datasets with stationary noise (mainly vertical stripes), the VSNR (Variational Stationary Noise remover) plugin in ImageJ was used to reduce the effect ^25^. The full datasets were segmented manually in IMOD ^26^, whereas some parts of the datasets were segmented semi-automatically in more detail using Ilastik ^27^ and MIB ^28^. For the full segmentation in Fig. 1a,b, every 10-15th slices was segmented and the slices in between were interpolated within IMOD. In the dataset of a cell at 11.2 min, a small part of the nucleus was missing and the NE in the region was reconstructed by interpolation. For the detailed segmentation in Fig. 1c, representative parts of the dataset, with the size of 5n10 μm in XY and 2 μm in Z of one set of daughter chromosomes, were selected and segmented every single slices for 3.1, 3.9, 4.3, 5.3, and 6.3 min and every two slices for 5.7 min. The datasets up to 4.3 min, where the ER is not yet firmly attached to the chromosomes, had to be manually segmented in MIB, since the automatic recognition of ER/chromosomes in Ilastik did not perform well. Datasets starting at 5.3 min, where the ER turns into the NE and firmly attaches to the chromosomes, were labeled in the pixel classification pipeline in Ilastik. The different objects were trained on a couple of slices of a dataset and the software predicted on other slices of the dataset. In the end, Ilastik created a simple segmentation file giving the highest probability for each pixel to be foreground (object). The remaining imprecise labelling was removed manually in MIB, and nuclear pores were added in IMOD. For Fig. 1c, the contrast of the FIB-SEM image was enhanced by projection of a series of 2–5 images for presentation purposes.

### Quantification of FIB-SEM data

For the analysis of chromosome coverage by the NE, the 3D models of fully-segmented chromosomes and ER were converted to lists of points every 10 pixels in IMOD. The minimal distance from each point on the chromosome surface to ER was measured in 3D space, and the ER within 100 nm away from chromosome surface was defined as the NE.

For measuring the size of membrane holes, segmented ER regions were connected in each Z-slice based on connected component analysis and location information. Each of the connected components in a slice was labeled first and its centroid was detected. A 2D matrix was generated that represents pairwise inter-points Euclidean distances of all centroids of ER regions. ER regions were then connected sequentially by utilizing this matrix in a customized breadth first search manner. Specifically, two ER regions having the shortest pairwise distance were selected in the first step. These two regions were connected by a straight line that bridges their nearest boundary pixels. In the second step, previous two regions were used as references to identify a third ER region that has the shortest distance from one of the two references. This region was connected to the closest reference region and taken as the second reference region for next step replacing the reference region it connects with. The connected network was grown gradually in similar fashion by selecting a new region with respect to the two references. One of the references was updated in each step and no region was connected more than twice. Thus, for n number of regions n-1 lines were generated that connects 2 × (n−1) boundary points of the ER regions. After having the list of line coordinates in each Z-slice, we connected lines of subsequent slices that were within proximity of one voxel, resulting in the 3D surface of the holes. The surface area was then quantified, and the diameter was calculated assuming circular shape for the intuitive understanding of the hole size in comparison to nuclear pores. The minimal distance measurement, connecting membranes and the surface area measurement were done in MATLAB (The MathWorks Inc., Natick, MA).

### Electron tomography

Electron tomography was performed as described in a previous report ^13^. Briefly, images were recorded over a −60° to 60° tilt range with an angular increment 1° at a pixel size of 0.75 nm with a TECNAI TF30 transmission EM (TEM; 300 kV; FEI, Hillsboro, OR) equipped with a 4k × 4k Eagle camera (FEI). Tomograms were reconstructed using R-weighted backprojection method implemented in the IMOD software package (version 4.6.40b) ^26^. For studying interphase NPCs, the cells at >3 h after anaphase onset were examined. For presentation purposes, the image contrast was enhanced by projections of 20 slices (corresponding to 15 nm) for Fig. 2a,b and **Supplementary Figs. 2a** and **3a**, side views in Fig. 4a, and side views in **Supplementary Fig. 8**, 8 slices (6 nm) for top views i and iii in Fig. 4a and all the top views in **Supplementary Fig. 8**, and 32 slices (24 nm) for top views ii in Fig. 4a. The surface area of the NE analyzed by EM tomography was measured, and the nuclear pore density was calculated as described previously ^13^. The overall structural similarity of the nuclear pores in freeze-substituted and heavy metal-stained cells to the respective cryo structures ^13^ indicates a good structure preservation and representation of actual molecular density of nuclear pores. Also negative staining analysis has been often done together with averaging and even molecular interpretation. Our interpretation relies on rather low resolution features such as the presence or absence of an entire ring, all of this is clearly apparent also prior to averaging (Figs. 2 and **4** and **Supplementary Fig. 8**).

### Membrane profile analysis

For Fig. 2b and **Supplementary Fig. 3**, the ER and nuclear membranes were manually traced and aligned as described in a previous report ^13^. The tip-to-tip distance, the ONM/INM distance, and the membrane tip curvature were measured from these two-dimensional profiles in MATLAB 7.4. For the ONM/INM distance, the median of the distance at 50 points between 45 and 90 nm away from the edge of nuclear pores was measured. For the tip curvature, the second derivative of the second-degree polynomial fit at each point along the profile was measured within an arc of 20 nm centered at the apex, and the inverse of the median value was shown as the radius of the tip curvature. The membrane holes of pre-pores are circle and rotationally symmetric in the plane parallel to the nuclear envelope (**Supplementary Fig. 8**), and therefore the membrane profile analyses done in 2D represent the actual structural features.

### Quantification of dense material within membrane holes

The tomographic subvolume around the membrane hole in the plane of the NE, as indicated by red dashed box in the left panel in Fig. 3a, was Z-projected. Then the radial intensity profile from the center towards the edge was measured using the radial profile plugin in ImageJ. The mean intensity in the region between 57 and 72 nm away from the center was defined as the intensity of the membrane lumen, and used to normalize the intensity in other regions for each membrane hole. The radial intensity profile was measured on 24 nm-Z-projected images for Fig. 3a and **Supplementary Fig. 8c**, and on 15 nm-Z-projected images for **Supplementary Fig. 3b**. For measuring inner ring intensity in **Supplementary Fig. 8c**, the region comprising 80% of the pore radius was regarded as inner ring, and the average intensity of the local 5 nm thick volume was quantified.

### Quantification of nuclear surface area growth

HeLa cells stably expressing histone H2b-mCherry were observed by confocal microscopy (LSM780; Carl Zeiss) using 63 × 1.4 NA Plan-Apochromat objective (Carl Zeiss) ^13^. Fluorescent chromatin was monitored under the following conditions: 40 optical sections, section thickness of 1.4 μm, z-stacks of every 0.7 μm, the xy resolution of 0.13 μm, and a time-lapse interval of 1 min. The chromosomes were segmented and the surface area was computed as described in a previous report ^13^. Fluorescence images were filtered with a median filter (kernel size: 0.25 × 0.25 μm) for presentation purposes. Visualization of the chromosome surface in 3D was done in MATLAB.

### Seriation of nuclear pores and particle averaging

The subtomograms containing individual pre- and mature pores were extracted and aligned without imposing any symmetry by using the previously described method ^29^. For seriation, the aligned 3D images of individual nuclear pores were resized from 160 × 160 × 120 pixels to 80 × 80 × 60 by averaging neighboring pixels. The whole data set of 399 nuclear pores was then represented by a tensor **A** of dimensions 80 × 80 × 60 × 399. We factorized this tensor into a core tensor **Z** and a set of 4 orthogonal basis matrices U_1_, U_2_, U_3_, and U_4_ using a truncated higher order singular value decomposition (HOSVD) ^30^,^31^ of rank (20,20,15,5):

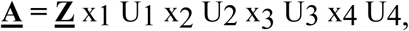

where xn represents the mode-n product. HOSVD can be thought of as a form of higher-order principal component analysis with the factor matrices Ui representing the principal components in each mode. U4 in particular can be seen as containing the coordinates of each nuclear pore on latent axes. From this, we computed the similarity between nuclear pores as Si,j = 1/(1+Di,j), where Di,j is the Euclidean distance between pore i and pore j in the space defined by the 5 components of U4.

Seriation aims at finding a good ordering of items such that similar items are close to each other in the sequence and dissimilar items are further apart or, equivalently, dissimilarities between items increases with their rank difference. This can be formalized using the 2-sum criterion which multiplies the similarity between items by the square of their rank difference. The problem is then to find the optimal ordering that globally minimizes this criterion:

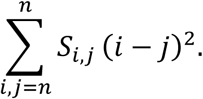

It has been shown that the optimal order for the 2-sum criterion can be recovered from the sorting order of the Fiedler vector (i.e. the second smallest eigenvector of the Laplacian of the similarity matrix) ^32,33^.

The ordered nuclear pores were sub-clustered into 5 classes with equal Fiedler vector value range (the Fiedler vector value of −1.0 – −0.6, −0.6 –-0.2, −0.2 – 0.2, 0.2 – 0.6, and 0.6 – 1.0 for the clusters 1−5, respectively). The original subtomograms (200 × 200 × 200 pixels) corresponding to the clustered pores were aligned and averaged in each cluster separately without imposing any symmetry by using the previously described method ^27^. For **Supplementary Fig. 8b**, pores in individual postmitotic times were selected, aligned, and averaged.

### Genome editing and southern blotting

Genome editing with CRISPR-Cas9 nickases was carried out as described in a previous report

^13^. The gRNA sequences for mEGFP-Nup205 are as follows: 5’AGAGCCCTTCTGCCGCATGGAGG3’ and 5’CTCTAAGATGGCGACGCCTTTGG3’ Southern blotting was performed as described in previous reports^34,35^ with the following modifications: Genomic DNA was prepared with the Wizard Genomic DNA Purification Kit (Promega, Madison, WI) and digested with *Nco*I and *Ssp*I (NEB, Ipswich, MA). The probe sequences are as follows: mEGFP-Nup205 (GCTACGAATTGAAGAAGCCTGTGAGAGAATGCTTGTAAAGTGTAATGATAGGTGAAGAGGAGTTGCTGGAATTTAAAGAGTTACTTTATAGAAAGCAAAGTTATCTACTGAGAGGGAAGCGTGTGGGCTTAAAAAGTGCATGTGGAGAGGGAAAGGCTTGTAATAGCTGTGCTGAGAGAGAAATCTTTGAATGCCGCTTCCTGTCAAGTTCCTGCTCAGTATTCTTCTCAGTTTCATGATAAACTTCTTGAAAAATTCCTTTGCGTTTACTTTCTCTCTTCCCCACTTCACTCCTTAAAACCACTTTAATCGTAGTTTGGCATTTGTCTC). mEGFP (CACATGAAGCAGCACGACTTCTTCAAGTCCGCCATGCCCGAAGGCTACGTCCAGGAGCGCACCATCTTCTTCAAGGACGACGGCAACTACAAGACCCGCGCCGAGGTGAAGTTCGAGGGCGACACCCTGGTGAACCGCATCGAGCTGAAGGGCATCGACTTCAAGGAGGACGGCAACATCCTGGGGCACAAGCTGGAGTACAACTACAACAGCCACAACGTCTATATCATGGCCGACAAGCAGAAGAACGGCATCAAGGTGAACTTCAAGATCCGCCACAACATCGAGGACGGCAGCGTGCAGCTCGCCGACCACTACCAGCAGAACACCC).

### Kinetic analysis of nucleoporin assembly in living cells

DNA was stained with 0.2 μM siliconnrhodamine Hoechst ^36^, and the nucleus and nucleoporins during mitosis exit were monitored by confocal microscopy (LSM780) using 63 × 1.4 NA Plan-Apochromat objective (Carl Zeiss) under the following conditions: 1 optical section, section thickness of 3.0 μm, the xy resolution of 0.13 μm, and a time-lapse interval of 12 sec. Total intensities of nucleoporins accumulating in the NE were quantified as described in a previous report ^6^. Fluorescence images were filtered with a median filter (kernel size: 0.25 × 0.25 μm) for presentation purposes.

### Sample size determination and statistical analysis

For FIB-SEM, we first obtained data from 3 different cells at 3.1, 5.3, and 6.3 min after AO as pilot experiments. After analyzing the chromosome coverage by the NE and nuclear pore number for these three cells, we decided to observe four more cells; three cells at 3.9, 4.3 and 5.7 min which are intermediate time points for the NE coverage and pore number increase, and one cell at 11.2 min for the end point (see Fig. 1). For EM tomography, we analysed 5−6 μm^2^ NE surface area in each cell at 4.8, 6.1, 10.2 and 15.0 min after AO and in interphase. After quantifying the pre-pore diameter, we analyzed one more cell at 7.7 min which is the intermediate time point for the diameter increase. Exact value of the analyzed surface area and the number of nuclear pores found are listed in **Supplementary Table 2**. We picked up all the membrane holes which were visible in the EM tomograms and less than 80 nm in diameter. Since for some pores the membrane contrast was too low to perform membrane profile analysis due to high noise, we excluded them form the quantitative structural analysis and used them only for measuring the pore density in Fig. 2d and **Supplementary Fig. 5c**. In addition, the nuclear pores near gold particles were removed from the spectral ordering, since the gold particles used for tomography alignment gave high electron density that interfered with the structure-based ordering of those pores. The exact number of pores excluded is also listed in **Supplementary Table 2**. For time-lapse imaging in **Supplementary Fig. 5a,b** and **Supplementary Fig. 10**, the data were from two independent experiments. Statistical analyses were performed only after all the data were taken. Statistical analysis methods, sample sizes (N) and P values (P) for each experiment are indicated in figure legends.

### Code availability

The computer codes used in this study are available on reasonable request from the corresponding author.

### Data availability

FIB-SEM datasets that support the finding of this study have been deposited to EMPIAR, the Electron Microscopy Public Image Archive (https://www.ebi.ac.uk/pdbe/emdb/empiar/; accession codes: 10100, 10101, 10102, 10103, 10104, 10105, 10109). Raw 2D-tilt series EM images for tomography are deposited to Electron Microscopy Data Bank (https://www.ebi.ac.uk/pdbe/emdb/; entry ID: EMD-3820). The other relevant datasets that are not included in the paper are available on request from the corresponding author.

## Acknowledgements

We thank the European Molecular Biology Laboratory Electron Microscopy Core Facility; the members of the Ellenberg group and the Beck group for advice and discussion; Anna Kreshuk and Ilya Belevich for help with using Ilastik and MIB, respectively. This work was supported by grants from the German Research Council to J.E. (DFG EL 246/3-2 within the priority program SPP1175), the Baden-Württemberg Stiftung to J.E., and the European Research Council to M.B. (309271-NPCAtlas), as well as by the European Molecular Biology Laboratory (S.O., A.M.S., M.S., J.K.H., M.J.H., S.S., Y.S., M.B., J.E.). S.O. was additionally supported by the EMBL Interdisciplinary Postdoc Programme (EIPOD) under Marie Curie Actions COFUND and a JSPS fellowship (The Japan Society for the Promotion of Science, postdoctoral fellowship for research abroad).

## Author contributions

SO, AMS, YS, MB, and JE designed the project. SO performed quantitative analysis of FIBSEM data and all the experiments and analyses of EM tomography. AMS acquired all the FIBSEM data and carried out segmentation of FIB-SEM images. MS contributed to the computational quantitative analysis of EM images. JKH performed spectral ordering. MJH carried out the segmentation of fluorescence images and assisted with computational analysis of EM images. SS helped with the segmentation of FIB-SEM images. MK generated genome-edited cell lines. YS, MB, and JE supervised the work. SO and JE wrote the paper. All authors contributed to the analysis and interpretation of data and provided input on the manuscript.

## References

1. Beck, M. & Hurt, E. The nuclear pore complex: understanding its function through structural insight. Nat. Rev. Mol. Cell Biol. 18, 73–89 (2017).

2. Ungricht, R. & Kutay, U. Mechanisms and functions of nuclear envelope remodelling. Nat. Rev. Mol. Cell Biol. 18, 229–245 (2017).

3. Weberruss, M. & Antonin, W. Perforating the nuclear boundary - how nuclear pore complexes assemble. J. Cell Sci. 129, 4439–4447 (2016).

4. LaJoie, D. & Ullman, K. S. Coordinated events of nuclear assembly. Curr. Opin. Cell Biol. 46, 39–45 (2017).

5. Dultz, E. et al. Systematic kinetic analysis of mitotic dis - and reassembly of the nuclear pore in living cells. J. Cell Biol. 180, 857–865 (2008).

6. Otsuka, S., Szymborska, A. & Ellenberg, J. Imaging the assembly, structure, and function of the nuclear pore inside cells. Methods Cell Biol. 122, 219–238 (2014).

7. Rabut, G., Lenart, P. & Ellenberg, J. Dynamics of nuclear pore complex organization through the cell cycle. Curr. Opin. Cell Biol. 16, 314–321 (2004).

8. Antonin, W., Ellenberg, J. & Dultz, E. Nuclear pore complex assembly through the cell cycle: regulation and membrane organization. FEBS lett. 582, 2004–2016 (2008).

9. Wandke, C. & Kutay, U. Enclosing chromatin: reassembly of the nucleus after open mitosis. Cell 152, 1222–1225 (2013).

10. Schellhaus, A. K., De Magistris, P. & Antonin, W. Nuclear Reformation at the End of Mitosis. J. Mol. Biol. (2015).

11. Fichtman, B., Ramos, C., Rasala, B., Harel, A. & Forbes, D. J. Inner/Outer nuclear membrane fusion in nuclear pore assembly: biochemical demonstration and molecular analysis. Mol. Biol. Cell 21, 4197–4211 (2010).

12. Lu, L., Ladinsky, M. S. & Kirchhausen, T. Formation of the postmitotic nuclear envelope from extended ER cisternae precedes nuclear pore assembly. J. Cell Biol. 194, 425–440 (2011).

13. Otsuka, S. et al. Nuclear pore assembly proceeds by an inside-out extrusion of the nuclear envelope. Elife 5 (2016)

14. Walther, T. C. et al. The conserved Nup107-160 complex is critical for nuclear pore complex assembly. Cell 113, 195–206 (2003).

15. Rotem, A. et al. Importin beta regulates the seeding of chromatin with initiation sites for nuclear pore assembly. Mol. Biol. Cell 20, 4031–4042 (2009).

16. Puhka, M., Joensuu, M., Vihinen, H., Belevich, I. & Jokitalo, E. Progressive sheet-totubule transformation is a general mechanism for endoplasmic reticulum partitioning in dividing mammalian cells. Mol. Biol. Cell 23, 2424–2432 (2012).

17. Goldberg, M. W., Wiese, C., Allen, T. D. & Wilson, K. L. Dimples, pores, star-rings, and thin rings on growing nuclear envelopes: evidence for structural intermediates in nuclear pore complex assembly. J. Cell Sci. 110, 409–420 (1997).

18. Kiseleva, E., Rutherford, S., Cotter, L. M., Allen, T. D. & Goldberg, M. W. Steps of nuclear pore complex disassembly and reassembly during mitosis in early Drosophila embryos. J. Cell Sci. 114, 3607–3618 (2001).

19. Kosinski, J. et al. Molecular architecture of the inner ring scaffold of the human nuclear pore complex. Science 352, 363–365 (2016).

20. Vollmer, B. et al. Dimerization and direct membrane interaction of Nup53 contribute to nuclear pore complex assembly. EMBO J. 31, 4072–4084 (2012).

21. Eisenhardt, N., Redolfi, J. & Antonin, W. Interaction of Nup53 with Ndc1 and Nup155 is required for nuclear pore complex assembly. J. Cell Sci. 127, 908–921 (2014).

22. Champion, L., Linder, M. I. & Kutay, U. Cellular Reorganization during Mitotic Entry. Trends Cell Biol. 27, 26–41 (2017).

23. Narayan, K. et al. Multi-resolution correlative focused ion beam scanning electron microscopy: applications to cell biology. J. Struct. Biol. 185, 278–284 (2014).

24. Cardona, A. et al. TrakEM2 software for neural circuit reconstruction. PLoS one 7, e38011 (2012).

25. Fehrenbach, J., Weiss, P. & Lorenzo, C. Variational algorithms to remove stationary noise: applications to microscopy imaging. IEEE transactions on image processing 21, 4420–4430 (2012).

26. Kremer, J. R., Mastronarde, D. N. & McIntosh, J. R. Computer visualization of three-dimensional image data using IMOD. J. Struct. Biol. 116, 71–76 (1996).

27. Sommer, C., Straehle, C., Koethe, U., Hamprecht, F. A. ilastik: Interactive Learning and Segmentation Toolkit. Proceedings of iEEE International Symposium on Biomedical Imaging (ISBI), 230–233 (2011).

28. Belevich, I., Joensuu, M., Kumar, D., Vihinen, H. & Jokitalo, E. Microscopy Image Browser: A Platform for Segmentation and Analysis of Multidimensional Datasets. PLoS Biol. 14, e1002340 (2016).

29. Beck, M. et al. Nuclear pore complex structure and dynamics revealed by cryoelectron tomography. Science 306, 1387–1390 (2004).

30. De Lathauwer, L., Moor, B. D. & Vandewalle, J. A. Multilinear Singular Value Decomposition. SIAM J. Matrix Anal. Appl. 21, 1253–1278 (2000).

31. Kolda, T. G. & Bader, B. W. Tensor Decompositions and Applications, SIAM Rev. 51, 455–500, (2009).

32. Atkins, J. E., Boman, E. G. & Hendrickson, B. A spectral algorithm for seriation and the consecutive ones problem. SIAM J. Computing 28, 297–310 (1998)

33. Ding, C. & He, X. Linearized cluster assignment via spectral ordering. Proceedings of the Twenty-first International Conference on Machine Learning (ICML '04) pp. 30(2004).

34. Mahen, R. et al. Comparative assessment of fluorescent transgene methods for quantitative imaging in human cells. Mol. Biol. Cell 25, 3610–3618, (2014).

35. Koch, B. et al. Generation and validation of homozygous fluorescent knock-in cells using genome editing. bioRxiv 188847 (2017).

36. Lukinavicius, G. et al. SiR-Hoechst is a far-red DNA stain for live-cell nanoscopy. Nat. Commun. 6, 8497 (2015)

37. Shimi, T., Butin -Israeli, V. & Goldman, R. D. The functions of the nuclear envelope in mediating the molecular crosstalk between the nucleus and the cytoplasm. Curr. Opin. Cell Biol. 24, 71–78 (2012).

